# Single-cell Transcriptomics Identify Extensive Heterogeneity in the Cellular Composition of Mouse Achilles Tendons

**DOI:** 10.1101/801266

**Authors:** Andrea J De Micheli, Jacob B Swanson, Nathaniel P Disser, Leandro M Martinez, Nicholas R Walker, David J Oliver, Benjamin D Cosgrove, Christopher L Mendias

**Author notes:** Corresponding Author Christopher Mendias, PhD, Hospital for Special Surgery, 535 E 70th St, New York, NY 10021, USA. These authors contributed equally to the manuscript (Andrea J De Micheli, Jacob B Swanson).

## Abstract

Tendon is a connective tissue that transmits forces between muscles and bones. Cellular heterogeneity is increasingly recognized as an important factor in the biological basis of tissue homeostasis and disease, but little is known about the diversity of cells that populate tendon. Our objective was to explore the heterogeneity of cells in mouse Achilles tendons using single-cell RNA sequencing. We assembled a transcriptomic atlas and identified 11 distinct cell types in tendons, including 3 previously undescribed populations of fibroblasts. Using trajectory inference analysis, we provide additional support for the notion that pericytes are progenitor cells for the fibroblasts that compose adult tendons. We also modeled cell-interactions and identified ligand-receptor pairs involved in tendon homeostasis. Our findings highlight notable heterogeneity between and within tendon cell populations, which may contribute to our understanding of tendon extracellular matrix assembly and maintenance, and inform the design of therapies to treat tendinopathies.

## Introduction

Tendon is a dense, collagen- and proteoglycan-rich connective tissue that transmits forces between muscles and bones. Tendon fibroblasts, or tenocytes, are thought to be responsible for extracellular matrix (ECM) production, organization, and maintenance **(Gumucio et al., 2015)**. During development and early postnatal stages, tendon is a relatively cellular tissue with high rates of cell proliferation, but 3 weeks after birth mice tendons become hypocellular with low rates of cellular turnover **(Grinstein et al., 2019; Mendias et al., 2012)**. Single-cell RNA sequencing (scRNAseq) is a technique that can expose the cellular heterogeneity within tissues. Cellular heterogeneity is increasingly recognized as important for the biological function of tissues, and for the development of new therapies for diseases **(Paolillo et al., 2019)**. Therefore, our objective was to determine the diversity in cell populations in postnatal tendons, and to establish an interactive transcriptomic atlas of tendon using scRNAseq as a resource to enable widespread exploration of the cellular heterogeneity of this functionally important connective tissue.

## Results and Discussion

We isolated Achilles tendons (N=4), the major load bearing tendons of the hindlimb, from 6-week old male C57BL/6J mice. This age was selected to be reflective of early adulthood, as the cells within tendon have low proliferation rates and are specified at this point, but the ECM is still actively being synthesized as the skeleton continues to slowly lengthen **(Somerville et al., 2004; Mendias et al., 2012; Grinstein et al., 2019)**. Tendons were digested to generate single-cell suspensions, and scRNAseq was performed to generate 4 independent single-cell datasets. Given the hypocellular nature of tendons, datasets were bioinformatically integrated to augment cell type coverage and improve gene expression analysis statistical power **(Stuart et al., 2019)** (Figure S1A). Collectively, we resolved 1197 cells after quality control validation, which clustered into 11 distinct populations (Figure 1A). Differences in cell type proportions between tendon samples likely reflect cell-sampling effects (Figure S1B). The top differentially expressed genes in each cell type are shown in Figure 1B. We published an online interactive atlas available at https://mendiaslab.shinyapps.io/Tendon_SingleCell_Atlas/. Flow cytometry using cell surface antigens was also performed (Figure 1E-F).

**Figure 1.**
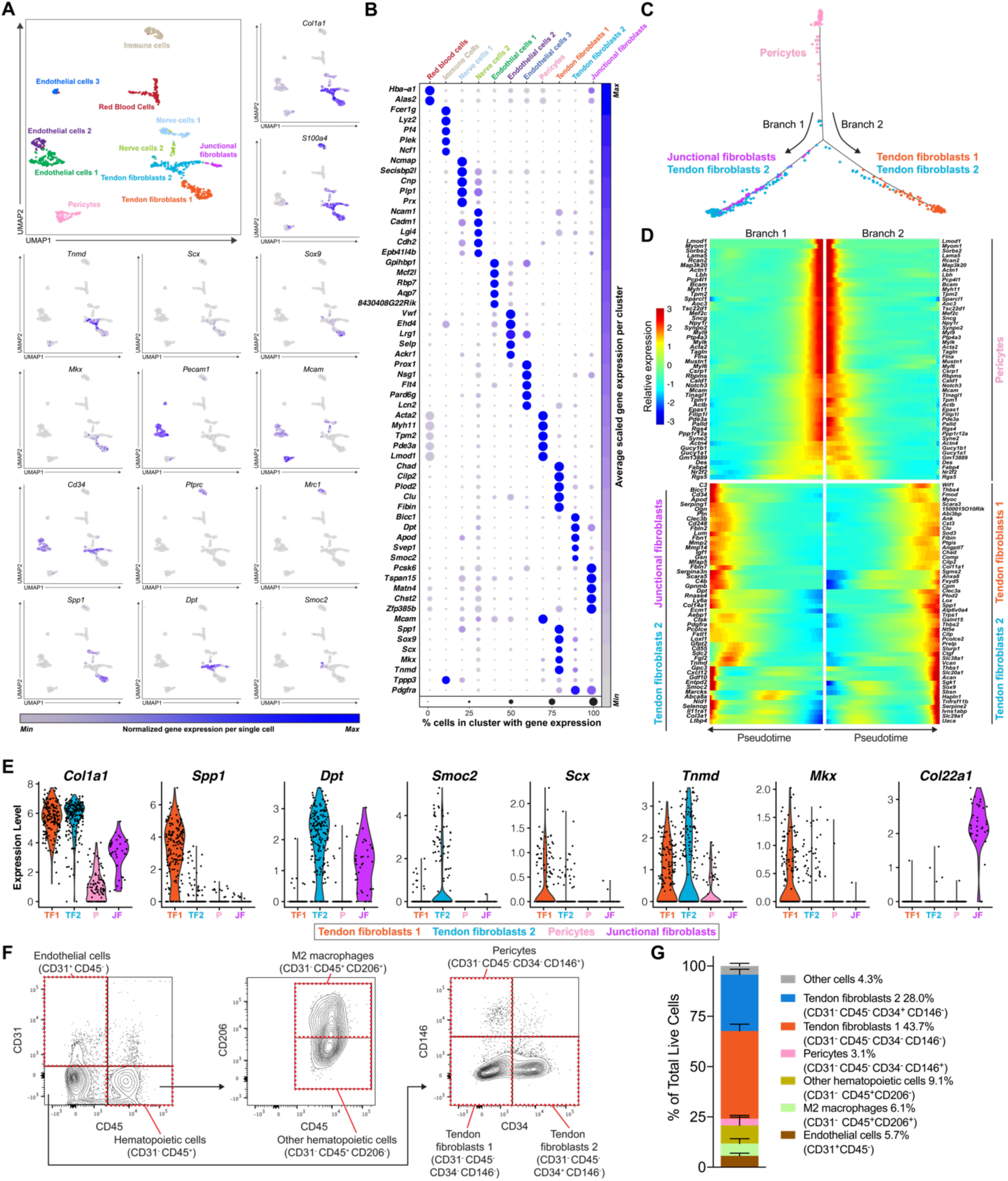
Single-cell transcriptomic atlas of the mouse Achilles tendon. (A) Single-cell transcriptomic atlas of the mouse Achilles tendons identifying 11 unique cell types. Expression of tendon gene sets mapped to the atlas. (B) Normalized expression of gene sets that are differentially enriched in each cell type. (C) Single-cell pseudotime trajectory of Pericytes into different Tendon fibroblast populations. (D) Heatmap demonstrating the top 50 genes in that are expressed across pseudotime progression from Pericytes to Tendon fibroblasts and Junctional fibroblasts subpopulations. (E) Violin plots for select tendon genes in Tendon fibroblasts (TF1, TF2), Junctional fibroblasts (JF), and Pericyte (P) subpopulations. (F) Flow cytometry contour plots and (G) cell quantification.

We identified three groups of fibroblasts that we refer to as tendon fibroblasts 1 and 2, and junctional fibroblasts (Figure 1A-B). These cells collectively express type I collagen (*Col1a1*) at a moderate to high level, and it is the basis of this *Col1a1* expression that we define these cells as fibroblasts (Figures 1A,B,E and 2A). One of these fibroblast groups displayed moderate *Col1a1* expression, and also expressed transcripts for type XXII collagen (*Col22a1*) at a high level. Type XXII collagen is known to be present at tissue junctions, and we therefore refer to these cells as junctional fibroblasts.

For flow cytometry, fibroblasts 1 are present in the CD31^-^CD45^-^CD34^-^CD146^-^ subpopulation, and fibroblasts 2 are in the CD31^-^CD45^-^CD34^+^CD146^-^ subpopulation (Figures 1F-G), while junctional fibroblasts are present in both. The fibroblast 1 and 2 subpopulations express somewhat distinct patterns of the ECM binding genes osteopontin (*Spp1*) and dermatopontin (*Dpt*), with osteopontin enriched in fibroblasts 1 and dermatopontin in fibroblasts 2 (Figures 1A,B,E). We therefore sought to verify the expression patterns of *Spp* and *Dpt* using RNA *in situ* hybridization (ISH). Similar to scRNAseq, *Spp1* and *Dpt* are expressed in separate cells (tendon fibroblasts 1 and 2, respectively), and occasionally in the same cell (Figure 2B). In addition, a subset of cells from the fibroblast 2 subpopulation also differentially express *Smoc2* (Figure 1A,E). Unlike RNA, at the protein level osteopontin, dermatopontin, and SMOC2 are mainly present in overlapping locations (Figure 2C). As tendon fibroblasts are arranged in linear clusters of several cells, these observations indicate that fibroblasts within clusters could be specialized to produce distinct proteins that constitute the ECM.

**Figure 2.**
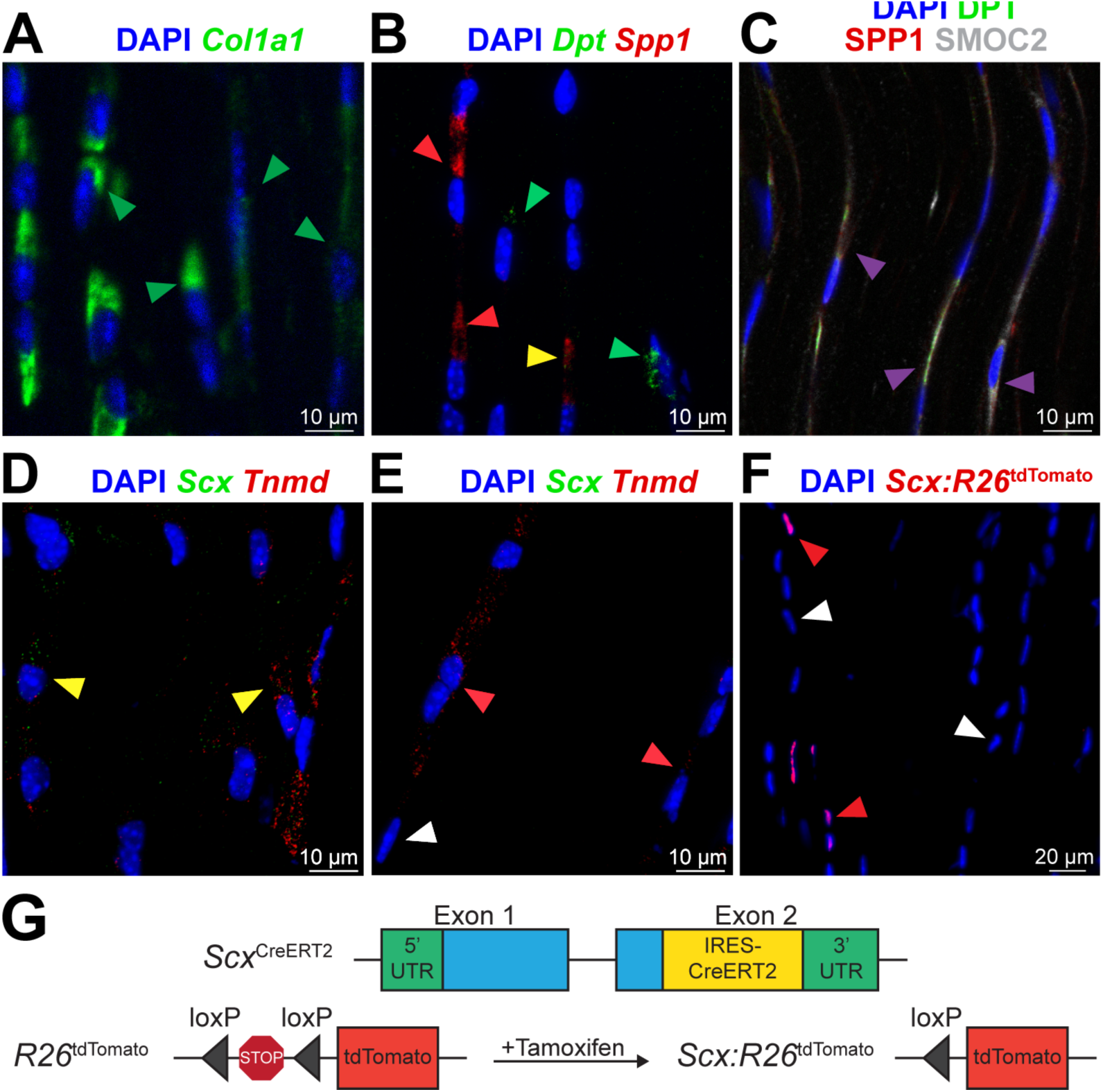
Histology. (A,B,D,E) RNA in situ hybridization and (C,F) protein fluorescence of Achilles tendons. (A) Col1a1 RNA, green; green arrows exemplify Col1a1+ fibroblasts. mRNA: Dpt, green; Spp1, red. Green arrows exemplify Spp1-Dpt+ fibroblasts; red arrows exemplify Spp1+Dpt-fibroblasts; yellow arrows exemplify Spp1+Dpt+ fibroblasts. Protein: DPT, green; osteopontin, red; and SMOC2, white. Magenta arrows exemplify overlap between DPT, SPP1, and SMOC2 protein. (D-E) RNA: Scx, green; Tnmd, red. Red arrows exemplify Scx-Tnmd+ fibroblasts; yellow arrows exemplify Scx+Tnmd+ fibroblasts; white arrows exemplify Scx-Tnmd- fibroblasts. (F) tdTomato protein identifies scleraxis-expressing cells (red arrows) compared to non-scleraxis expressing cells (white arrows). Nuclei visualized with DAPI. (G) Genetics of scleraxis lineage tracing mice.

Multiple studies have attempted to identify markers of the tenogenic lineage **(Bi et al., 2007).** Scleraxis (*Scx*) marks cells in the tenogenic lineage and is required for proper embryonic tendon development **(Huang et al., 2015)**. Several studies have used a transgenic reporter mouse to identify *Scx*-expressing cells (ScxGFP). ScxGFP mice express *GFP* under the control of a 4kb upstream regulatory sequence of *Scx*, but not the full length, endogenous *Scx* locus **(Pryce et al., 2007)**. Through 2 months of age, nearly all tendon fibroblasts of ScxGFP mice robustly express GFP, resulting in the widespread assumption that scleraxis is present in nearly every tendon fibroblast at this age. In addition to scleraxis, tenomodulin (*Tnmd*) is a transmembrane protein and mohawk (*Mkx*) is a transcription factor that are thought to also be consistent markers of the tenogenic lineage **(Huang et al., 2015)**. We therefore expected to observe widespread *Scx, Tnmd*, and *Mkx* expression in fibroblasts. To our surprise, a minority of fibroblasts 1 express *Scx*, and a portion of fibroblasts 1 and 2 express *Tnmd* (Figures 1A,B,E). *Mkx* was also only expressed in a subset of fibroblasts 1 and 2, with a greater proportion of fibroblasts 1 expressing *Mkx* than fibroblasts 2. We performed RNA ISH for *Scx* and *Tnmd* to confirm these findings. Consistent with scRNAseq, while all of the fibroblasts express *Col1a1*, only a subset express *Scx* and *Tnmd* (Figure 2A,D). Some cells expressed *Tnmd* but not *Scx* (Figure 2E). We also confirmed these findings with scleraxis lineage tracing mice, in which *CreERT2* is driven from the native *Scx* locus and a flox-stop-flox-tdTomato sequence is expressed from the ubiquitous *Rosa26* locus. This allows for *Scx-*expressing cells to permanently express *tdTomato* upon treatment with tamoxifen, referred to as *Scx*:*R26*^*tdTomato*^ mice (Figure 2G). Following 5 days of tamoxifen treatment, only a small portion of fibroblasts of *Scx*:*R26*^*tdTomato*^ mice contained tdTomato (Figure 2F), providing further support that *Scx* is only expressed in a subset of adult tendon fibroblasts. As *ScxGFP* contains only portions of the regulatory elements of native *Scx*, it is likely that ScxGFP mice overestimate *Scx* expression in adult tendons. While *Col1a1* is expressed in a small portion of endothelial and nerve cells, *Col1a1* may be more useful than *Scx* in lineage tracing or conditional deletion studies across all fibroblasts in tendons.

Pericytes, which express the transmembrane protein CD146 (*Mcam*), are thought to be a progenitor cell population for adult tendon fibroblasts **(Lee et al., 2015)**. To further explore this relationship, we used the cellular trajectory inference algorithm Monocle **(Qiu et al., 2017)** to model pericyte fate decisions in pseudotime. In support of a role for pericytes as progenitor cells, Monocle predicted a differentiation trajectory of pericytes into two branches consisting of either junctional fibroblasts and fibroblasts 2, or fibroblasts 1 and 2 (Figure 1C,D). The tubulin polymerization-promoting protein (*Tppp3*) has also been proposed as a marker of tendon progenitors as *Tppp3*^+^ *Pdgfra*^+^ cells **(Harvey et al., 2019)**. We found that *Tppp3* is differentially expressed by the tissue-resident immune subpopulation (log2fc = 2.5; q-value = 1.4e-28; Table S1) and is co-expressed at low levels with *Pdgfra* in a subset of fibroblasts (Figure 1B, S2). These cells also express *Col1a1* and therefore likely do not function entirely as progenitor cells. Previous scRNAseq reported abundant osteoblasts found in patellar tendons **(Harvey et al., 2019)**, however we observed only a small amount of cells expressing canonical osteoblast markers like *Runx2* (Figure S2B), with no detection of osteoblasts markers bone sialoprotein (*Bsp*), *Msx2*, or osterix (*Osx*) **(Rutkovskiy et al., 2016)**. This is consistent with whole tissue bulk RNAseq of Achilles, forepaw flexor, patellar, and supraspinatus tendons of mice and rats **(Disser et al., 2020)**, and together suggests a low abundance of osteoblasts in tendons.

Several other cell types were identified with scRNAseq (Figure 1A-B), and select subpopulations were quantified with flow cytometry (Figure 1F,G). *Sox9* is expressed in a portion of cells in the two fibroblast subpopulations, some of which express *Scx*, which is consistent with *Scx*^+^/*Sox9*^+^ and *Scx*^-^/*Sox9*^+^ cells previously reported in tendon **(Huang et al., 2015)**. Three clusters of endothelial cells were identified based on CD31 (*Pecam1*) expression and absence of the hematopoietic lineage marker CD45 (*Ptprc*), and quantified in flow cytometry as CD31^+^CD45^-^ cells. These three clusters are likely cells with specialized blood or lymphatic vascular functions. Red blood cells were identified by hemoglobin expression (*Hba-a1*). CD45 demarcated immune cells, with most expressing the tissue resident M2 macrophage marker CD206 (*Mrc1*) **(Schwartz et al., 2015)**. These cells were detected in flow cytometry as CD31^-^CD45^+^CD206^-^ cells for other hematopoietic cells, and CD31^-^CD45^+^CD206^+^ cells for M2 macrophages. We also identified two subpopulations of nerve cells. Nerve cells 1 expressed high levels of myelin proteolipid protein (*Plp1*) and Periaxin (*Prx*). Nerve cells 2 expressed high levels of neural cell adhesion molecule (*Ncam1*) and *Lgi4*. Nerve cells 1 are likely Schwann cells which surround the afferent sensory axons that innervate the sensory nerve structures of tendons. Nerve cells 2 could be glial cells that support the structural and metabolic activities of Schwann cells **(Nishino et al., 2010)**.

We then used a ligand-receptor model to chart scRNAseq data cell-interactions within the tendon tissue. Our model suggests >100 interactions between receptors that are differentially expressed in fibroblasts and ligands expressed by other cell types. Most of these ligands were highly expressed by immune cells or pericytes (Figure 3A-C). Among the top 50 immune-fibroblast or pericyte-fibroblast interactions, we identified TGFβ family ligands and receptors. TGFβ signaling is required for the maintenance of tendon progenitors and are strong inducers of tenogenic markers such as *Scx* **(Havis et al., 2014; Tan et al., 2020)**. Our results suggested that immune cells might be an important source of TGFβ signaling in tendon acting through CD109 and SDC2 receptors expressed by fibroblasts (Figure 3A). Furthermore, our model also identified that connective tissue growth factor (*Ctgf)* is differentially expressed by junctional and fibroblast 1 cells (but not 2) and interacts with at least 5 different receptors (*Lrp1, Lrp6, Itga5, Itgb2*, and *Itgam*). CTGF is recognized for its pro-tenogenic potential and its receptors on tendon progenitor cells may be a therapeutic target for tendon injuries **(Lee et al., 2015)**. CTGF interacts with LRP1, which has been documented to regulate extracellular remodeling **(Gaultier et al., 2010)**, is differentially expressed by all 3 fibroblast subpopulations (Figure 3D). LRP1 is also involved in a multitude of stem cell processes, such as neurogenesis, adipogenesis, and vascular development **(Masson et al., 2009; Nakajima et al., 2014; Safina et al., 2016)**, but LRP1 signaling in tenocyte differentiation has not been elucidated.

**Figure 3.**
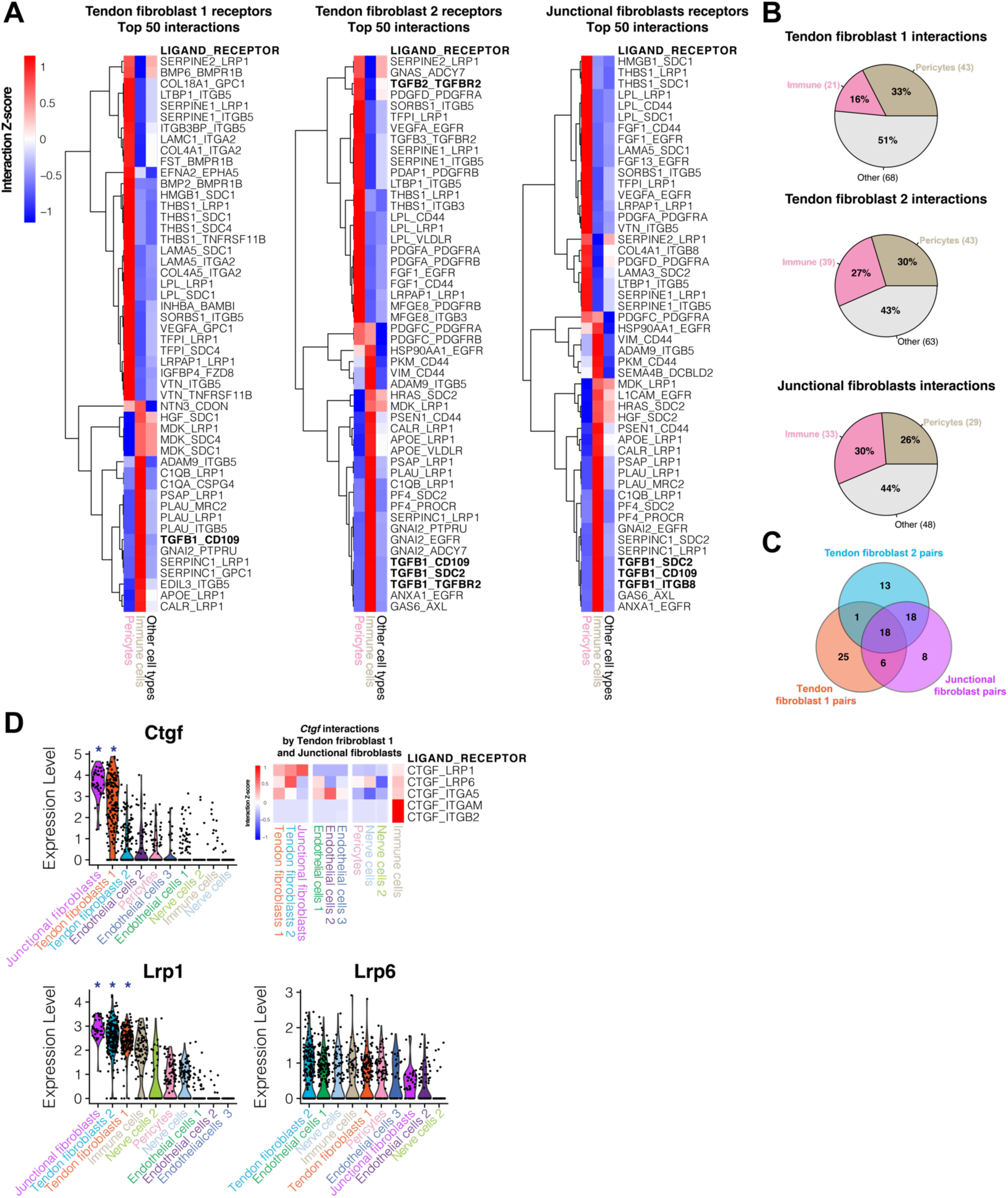
Ligand-receptor interaction model. (A) Heatmap of row-normalized (Z-score) ligand-receptor interactions scores. Scores are calculated from receptors differentially expressed in Fibroblasts subpopulations (Tendon fibroblast 1, 2 and Junctional compared other cell types) and ligands expressed by Pericytes and Immune cells or other cell types. Only the top 50 interactions ranked by score enriched in Pericyte and Immune cells are displayed. (B) Percentage of total interactions detected by the model enriched in Pericytes, Immune cells, or other cell types. (C) Venn diagram representing the total interactions pairs shared between Fibroblast subpopulations. (D) Violin plots for ligand-receptor genes ranked by average expression value. An asterisk indicated the gene is differentially expressed with p-value < 0.05. Heatmap of row-normalized ligand-receptor interactions scores involving the *CTGF* ligand differentially expressed by the Tendon fibroblast 1 and Junctional fibroblast subpopulations and receptors identified by the model that are expressed by other cell types.

There are several limitations to this study. We used a single age and sex of mice, and while we do not know how cell subpopulations will change with aging, we expect our findings are relevant to both sexes. There were fewer total cells analyzed for scRNAseq than other tissues **(Schaum et al., 2018)**, but this is likely due to the hypocellular nature of tendons. Despite these limitations, we provided datasets from 4 individual mouse tendons compared to other studies that pooled multiple tendon tissues from a single mouse **(Harvey et al., 2019; Tan et al., 2020)**, and our overall findings reveal tremendous heterogeneity in the cells that populate tendons. We identify 3 previously undescribed subpopulations of tendon fibroblasts based on relatively distinct expression of ECM proteins, and support the notion that pericytes are progenitor cells of adult fibroblasts. We also modeled ligand-receptor interactions in tendon that confirmed prior observations about TGFβ and CTGF, while also proposing new signaling targets. Our findings also indicated that fibroblasts could specialize to produce distinct components of the tendon ECM, which may have important implications in the treatment of tendinopathies.

## Materials and Methods

### Animals

This study was approved by the Hospital for Special Surgery/Weill Cornell Medical College/Memorial Sloan Kettering Cancer Center IACUC. Male 6-week old C57BL/6J mice (strain 000664) were obtained from the Jackson Laboratory (Bar Harbor, ME, USA). *Scx* ^*CreERT2*^ mice in which an IRES-CreERT2 sequence was inserted between the stop codon and 3’ UTR in exon 2 of scleraxis **(Howell et al., 2017)**, were kindly provided by Dr. Ronen Schweitzer (Shriners Hospitals for Children, Portland, OR, USA). We also obtained *R26*tdTomato reporter mice **(Madisen et al., 2010)** in which the constitutively expressed *Rosa26* locus was modified to contain a stop codon cassette flanked by loxP sites upstream of the red fluorescent tdTomato gene (Jackson Labs strain 007909). *Scx*^*CreERT2*^ mice were crossed to *R26* ^*tdTomato*^ mice and backcrossed again until homozygosity for both alleles was achieved. Six week old male *Scx*^*CreERT2/CreERT2*^:*R26*^*tdTomato/tdTomato*^ mice received an intraperitoneal injection of 1mg of tamoxifen (Millipore Sigma, St. Louis, MO, USA) dissolved in 50μL of corn oil for a period of 5 days as described **(Disser et al., 2019)** to induce recombination at the *R26* locus in scleraxis-expressing cells, generating animals referred to as *Scx*:*R26* ^*tdTomato*^ mice (Figure 2G).

### Surgical procedure

Mice were euthanized by exposure to CO2 followed by cervical dislocation. To remove Achilles tendons, a longitudinal incision through the skin was made down the midline of the posterior aspect of the lower limb, superficial to the gastrocnemius and Achilles tendon. The paratenon was reflected, and a sharp transverse incision was made just distal to the myotendinous junction and again just proximal to the enthesis, and the Achilles tendon was carefully removed. The procedure was performed bilaterally.

### Single cell isolation

The Achilles tendons were digested to obtain a single cell suspension, as modified from a previous study **(De Micheli et al., 2020)**. Two Achilles tendons from each animal were processed together as a single sample, and the tendons of N=4 mice were used. Tendons were finely minced using a scalpel, and then digested for 1 h at 37°C in a vigorously shaking solution consisting of 16 mg of Collagenase D (Roche, Pleasanton, CA, USA), 3.0 U of Dispase II (Roche), 640 ng of DNase I (Millipore Sigma, St. Louis, MO, USA), 20 μL of 4% bovine serum albumin (BSA, Millipore Sigma) in 2mL of low glucose DMEM (Corning, Corning, NY, USA). After digestion, the single cell suspension was filtered for debris using a 70 μm cell strainer and resuspended in 0.04% BSA in PBS.

### Single cell RNA sequencing and analysis

Single cell RNA sequencing and analysis was performed, as modified from a previous study **(De Micheli et al., 2020)**. Libraries were prepared using a Chromium Single Cell 3’ Reagent Kit (version 3, 10X Genomics, Pleasanton, CA, USA) following the directions of the manufacturer. Cells were loaded into the kit and processed using a Chromium Controller (10X Genomics). Following library preparation and quantification, libraries were sequenced by the Weill Cornell Medical College Epigenomics Core using a HiSeq 2500 system (Illumina, San Diego, CA, USA) Libraries were sequenced to generate approximately 250 million reads per sample, which resulted in on average approximately 960,000 reads per cell. Sequencing data has been deposited to NIH GEO (ascension GSE138515). Gene expression matrices were generated from the sequencing data using Cell Ranger (version 3.0.1, 10X Genomics) and the mm10 reference genome. Downstream analyses were carried out with R version 3.5.2 (2018-12-20) and the Seurat 3.1.0 R package **(Stuart et al., 2019)**. We integrated the four gene expression matrices for more powerful statistical analyses using SCTransform of Seurat method and evaluated differences in population number across samples (Figure S1). Genes expressed in less than 3 cells as well as cells <1000 unique molecular identifiers (UMIs) and <200 genes were removed from the gene expression matrix. We also filtered out cells with >20% UMIs mapping to mitochondrial genes. A total of 1197 cells remained after applying these criteria. We performed principal component analysis (PCA) and used the first 15 PCs for population clustering (unsupervised shared nearest neighbor, SNN, resolution=0.4) and data visualization (UMAP). Finally, differential expression analysis was achieved using the “FindAllMarkers” function in Seurat using a likelihood test that assumes a negative binomial distribution (min log2 fold-change > 0.25, min fraction > 25%). We refer to normalized gene expression values as the number of log-normalized counts per gene relative to the total number of counts per cell. Supplemental Table S1 provides differential gene expression data for each cell presented in the atlas and dot plot (Figure 1A-B).

### Single cell trajectory analysis

We used the Monocle v. 2.8.0 R package **(Qiu et al., 2017)** to infer a hierarchical organization between subpopulations of pericytes, fibroblasts 1 and 2, and junctional fibroblasts, to organize these cells in pseudotime. We took these subpopulations from the Seurat dataset from which we reperformed SNN clustering and differential expression analysis as described above. We then selected the top 300 differentially expressed genes based on fold-change expression with a minimum adjusted p-value of 0.01 for Monocle to order the cells using the DDRTree method and reverse graph embedding. We then used branch expression analysis to identify branch-dependent differentially expressed genes.

### Ligand-receptor interaction model

The model aims at scoring potential ligand-receptor interactions between fibroblast subpopulations and other cell types. We used the ligand-receptor interaction database from Skelly et al. **(Skelly et al., 2018)**. To calculate the score for a given ligand-receptor pair, we multiply the average receptor expression in fibroblasts with the average ligand expression per other cell type (including other fibroblast subpopulations to consider autocrine interactions). We only considered receptors that are differentially expressed in either of the 3 fibroblasts subpopulations.

### Flow cytometry

Tendons of N=3 mice were digested to obtain a single cell suspension as described above. Cells were treated with FC Block (BD, San Diego, CA, USA) and labeled with antibodies against CD31 (FAB3628N, R&D Systems, Minneapolis, MN, USA), CD34 (551387, BD), CD45 (103115, BioLegend, San Diego, CA, USA), CD146 (134707, BioLegend), and CD206 (141711, BioLegend), as well as DAPI (Millipore Sigma). Cytometry was performed using a FACSCanto (BD) and FlowJo software (version 10, BD). Forward Scatter (FSC) and Side Scatter (SSC) plots were used to identify cell populations. FSC-A (area) and FSC-H (height) were used to exclude doublets. Dead cells were excluded by DAPI signal. Cell populations are expressed as a percentage of total viable cells.

### RNA in situ hybridization

RNA in situ hybridization (ISH) was performed using a RNAscope HiPlex kit (ACD Bio-Techne, Newark, CA, USA) and detection probes against *Col1a1, Spp1, Dpt, Scx*, and *Tnmd* following directions of the manufacturer. Mouse Achilles tendons (N=4) for RNA ISH were snap frozen in Tissue-Tek OCT Compound (Sakura Finetek, Torrance, CA, USA) and stored at −80°C until use. Longitudinal 10 μm sections of tendons were prepared using a cryostat, and tissue sections were digested with protease, hybridized with target probes, amplified, and labeled with fluorophores. Tissue was counterstained with DAPI to visualize nuclei, and slides were imaged with the LSM 880 confocal microscope (Zeiss, Thornwood, NY, USA).

### Histology

Histology was performed as described **(Schwartz et al., 2015; Sugg et al., 2018).** Longitudinal sections of tendons were fixed with 4% paraformaldehyde, permeabilized with 0.1% Triton X-100. For antibody immunofluorescence (N=4), sections were blocked with a Mouse on Mouse Blocking Kit (Vector Labs, Burlingame, CA, USA) and incubated with primary antibodies against osteopontin (NB100-1883, Novus Biologicals, Centennial, CO, USA), dermatopontin (10537, Proteintech, Rosemont, IL, USA), and SMOC2 (sc-376104, Santa Cruz Biotechnology, Santa Cruz, CA, USA). Secondary antibodies conjugated to AlexaFluor 488, 555, and 647 (Thermo Fisher) were used to detect primary antibodies. For *Scx*:*R26*^*tdTomato*^ fluorescent imaging (N=4). Nuclei were identified in both sets of experiments with DAPI. Slides were visualized as described above.

## Acknowledgements

This work was supported by NIH grants R01-AR063649 (CLM) and R01-AG058630 (BDC), the Tow Foundation for the David Z Rosensweig Genomics Center (CLM), and by a Glenn Medical Research Foundation and American Federation for Aging Research Grant for Junior Faculty (BDC). We would like to thank Jonathan Daley, Richard Lee, and Marc Strum from the Hospital for Special Surgery for assistance in preparing the online single-cell atlas.

## Figures

**Figure S1.**
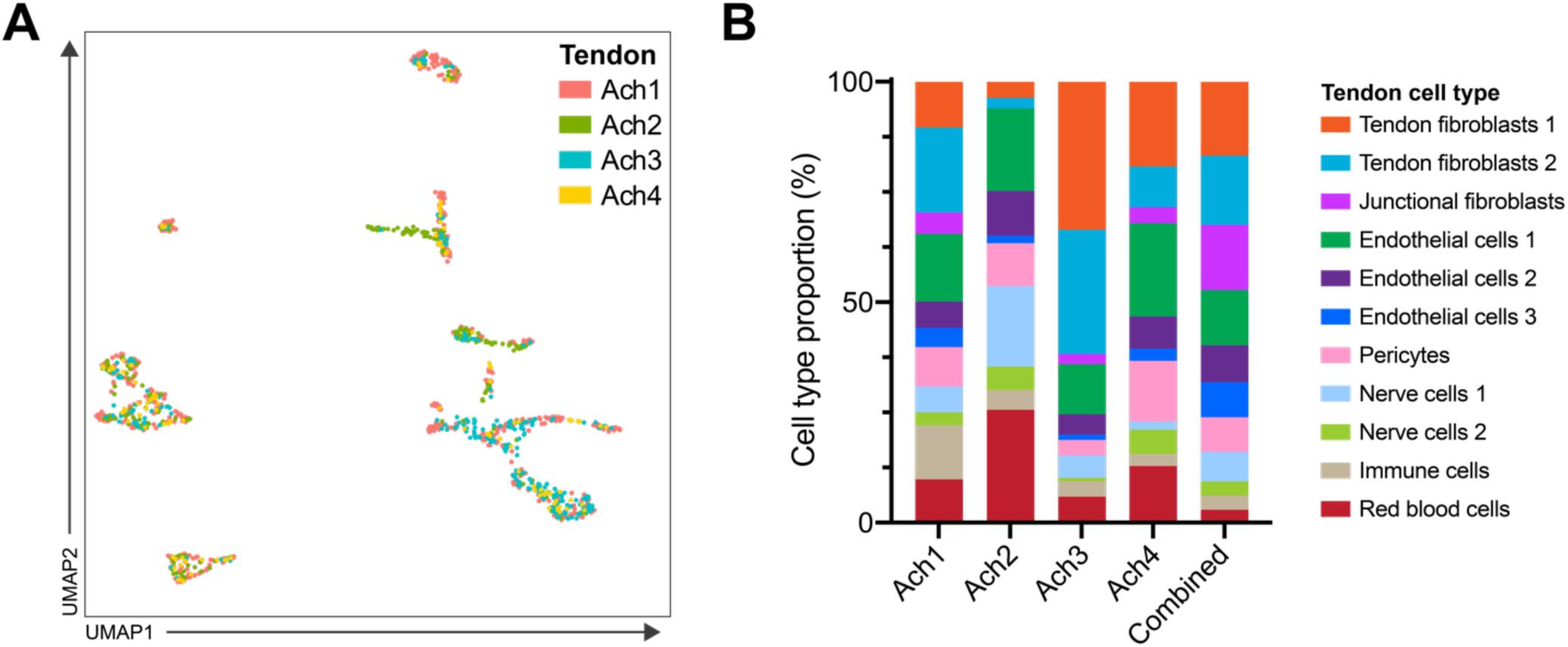
Bioinformatic integration of 4 mouse Achilles tendon scRNAseq datasets. (A) Single-cell transcriptomic atlas as in Figure 1A colored by Achilles (Ach) tendon sample. (B) Differences in cell type proportions across the 4 Ach tendon samples.

**Figure S2.**
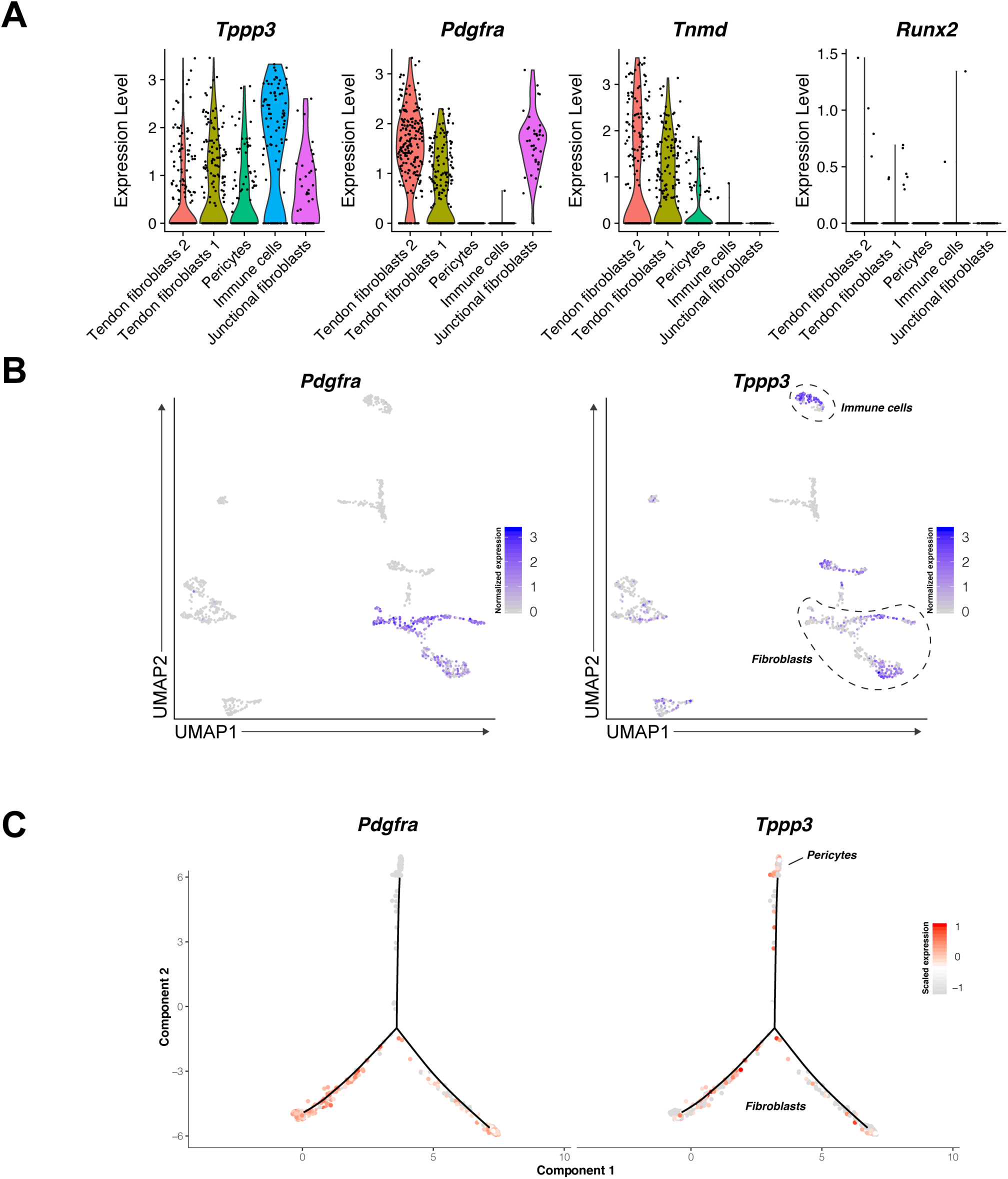
Pdgfra, Tppp3, Tnmd, and Runx2 expression from scRNAseq atlas. (A) Violin plots with normalized expression levels for select subpopulations. Expression of other osteoblast markers (*Bsp, Msx2, Osx*) is zero for all cells. (B) Whole atlas feature plot with normalized gene expression values. (C) Log Z-scaled relative expression along Monocle trajectory.

